# Photosymbiont density is correlated with constitutive and induced immunity in the facultatively symbiotic coral, *Astrangia poculata*

**DOI:** 10.1101/2024.02.28.582547

**Authors:** Isabella V. Changsut, Erin M. Borbee, Haley R. Womack, Alicia Shickle, Koty H. Sharp, Lauren E. Fuess

## Abstract

Scleractinian corals, essential ecosystem engineers that form the base of coral reef ecosystems, have faced unprecedented mortality in recent decades due climate-change related stressors, including disease outbreaks. Despite this emergent threat to corals, many questions still remain regarding mechanisms underlying observed variation in disease susceptibility. Emergent data suggests at least some degree of variation in disease response may be linked to variability in the relationship between host corals and their algal photosymbionts (Family Symbiodineaceae). Still, the nuances of connections between symbiosis and immunity in cnidarians, including scleractinian corals, remain poorly understood. Here we leveraged an emergent model species, the facultatively symbiotic, temperate, scleractinian coral *Astrangia poculata*, to investigate associations between symbiont density and both constitutive and induced immunity. We used a combination of controlled immune challenges with heat-inactivated pathogens and transcriptomic analyses. Our results demonstrate that *A. poculata* mounts a robust initial response to pathogenic stimuli that is highly similar to responses documented in tropical corals. Furthermore, we document positive associations between symbiont density and both constitutive and induced immune responses, in agreement with recent preliminary studies in *A. poculata*. A suite of immune genes, including those coding for antioxidant peroxidoxin biosynthesis, are constitutively positively associated with symbiont density in *A. poculata*. Furthermore, variation in symbiont density is associated with distinct patterns of immune response; low symbiont density corals induce preventative immune mechanisms whereas high symbiont density corals mobilize energetic resources to fuel humoral immune responses. In summary, our study reveals the need for more nuanced study of symbiosis-immune interplay across diverse scleractinian corals, preferably including quantitative energy budget analysis for full disentanglement of these complex associations and their effects on pathogen susceptibility.

## INTRODUCTION

Corals form the structural basis of coral reef communities and are essential in maintaining immense biodiversity and contributing to numerous ecosystem services (Alvarez-Filip et al. 2009; Eddy et al. 2021). However, these ecosystems have recently experienced significant degradation due to coral mortality, often linked to anthropogenically-driven climate change (Hoegh-Guldberg 1999; Wild et al. 2011). Indeed, global coral populations have experienced significant mortality as the result of temperature-related bleaching (the loss of the photosymbionts upon which corals depend) and outbreaks of pathogenic disease (Cramer et al. 2020; Croquer and Weil 2009; Estrada-Saldívar et al. 2020). Corals are susceptible to a wide range of pathogens; to date approximately 20 types of coral disease have been described (Green and Bruckner 2000; Morais et al. 2022; Weil et al. 2006). Interestingly, susceptibility to such diseases is highly heterogeneous both within and among species (Alvarez-Filip et al. 2019; Weil et al. 2006). The widespread effects of coral disease on global populations necessitates an improved understanding of coral immunity and the factors driving variability in disease outcomes within and between species.

Despite their position as basal metazoans, corals have a highly complex array of innate immune defenses (Bosch 2013; Mydlarz et al. 2016; Palmer and Traylor-Knowles 2012). For example, cnidarians possess some of the greatest diversity in pattern recognition receptors (PRRs) amongst all metazoans (Emery et al. 2021; Parisi et al. 2020; Zarate-Potes et al. 2019). These PRRs are responsible for identification of a wide diversity of both pathogenic and commensal microbiota, contributing to the maintenance of a healthy coral “holobiont” (host coral + commensal microbes; (Emery et al. 2021; Palmer and Traylor-Knowles 2012; Poole and Weis 2014; Weis 2019). Upon recognition of pathogenic bacteria, these PRRs induce signaling cascades which trigger effector responses including the production of diverse antimicrobial peptides (Mydlarz et al. 2016; Vidal-Dupiol et al. 2011), cytotoxic and encapsulating melanin (Mydlarz and Palmer 2011; Palmer et al. 2008), and antioxidants (Mydlarz and Harvell 2007). Notably, the cnidarian immune system, and particularly PRRs, also play important roles in the regulation of the diverse symbiotic associations which corals host (Bove et al. 2022; Weis 2019).

Corals form associations with diverse microbes including bacteria (Rohwer et al. 2002), viruses (Bettarel et al. 2015; Thurber and Correa 2011), and fungi (Amend et al. 2012; Raghukumar and Ravindran 2012). However, one of the most significant of coral symbiotic relationships is that between the coral host and photosymbiotic algae of the family Symbiodiniaceae (LaJeunesse et al. 2018; Simpson et al. 2011; van Oppen and Medina 2020). These photosymbionts are paramount to the success of the holobiont, providing essential inorganic nutrients to the host (Kopp et al. 2013; Wiedenmann et al. 2023). Associations between corals and Symbiodiniaceae are exceptionally diverse, demonstrating immense natural variation within and between species (LaJeunesse et al. 2018). This variability in symbiotic associations has been hypothesized to contribute to coral response to heat stress and bleaching, among other abiotic stressors (Baker 2004; Day et al. 2008; Kitchen et al. 2022). Furthermore, as the immune system is essential in regulating all stages of coral-Symbiodiniaceae relationships (Davy et al. 2012; Weis et al. 2001; Weis et al. 2008), it is hypothesized that this variability in coral-Symbiodiniaceae relationships affects pathogen response. A growing body of evidence supports a link between variation in coral symbioses and host immunity though the nuances of this relationship remain unclear (Changsut et al. 2022; Fuess et al. 2020b; Klein et al. 2024; Rivera and Davies 2021; Villafranca et al. 2023). Further complicating the understanding of this association, the vast majority of studies exploring this association have been conducted using obligately symbiotic coral species, which display limited variation in symbiont density. While several studies of both obligately and facultatively symbiotic corals have indicated a negative association between symbiont density and immunity (Fuess et al. 2020b; Rivera and Davies 2021), recent investigation of these same patterns in other species of facultatively symbiotic corals have revealed the opposite pattern (Changsut et al. 2022; Harman et al. 2022; Villafranca et al. 2023). These discrepancies highlight the need for broad investigation of the associations between symbiosis and immunity in cnidarians to further clarify these relationships.

Here we used the facultatively symbiotic scleractinian coral species *Astrangia poculata* to identify mechanisms underlying the links between symbiosis and pathogen response in cnidarians. Experimental work with *A. poculata*, which displays immense natural variation in symbiont density, enables hypothesis testing regarding symbiosis-immune interplay in cnidarians. Using a combination of experimental immune challenge and transcriptomic analyses, we characterized both the general immune response of *A. poculata* and the effects of symbiont density on immunity.

## METHODS

### Study System

*Astrangia poculata*, a temperate, facultatively symbiotic coral, is an emerging model system for addressing questions related to coral immunology and symbiosis. Unlike many tropical corals, *Astrangia poculata* is of a healthy IUCN status and can be collected and maintained in aquaria with relative ease. Colonies of *A. poculata* demonstrate extreme natural variation in density of their photosymbiont *Breviolum psygmophilum*, allowing for the study of symbiont density variation while avoiding the use of an external experimental stressor (Sharp et al. 2017). Symbiotic states can be roughly classified based on color, which typically correlates with *B. psygmophilum* density (“brown” >10^6^cells cm^-2^, “white” 10^4^ -10^6^ cm^-2^; (Sharp et al. 2017)).

### Sample Collection and Husbandry

Entire colonies of “white” (low symbiont density; *n* = 26) and “brown” (high symbiont densities; *n* = 11) *A. poculata* were collected from Fort Wetherill in Jamestown, Rhode Island (41°28′40″ N, 71°21′34″ W) in June of 2021, at a depth of 5-15 meters. Colonies were shipped to Texas State University where they were maintained in ten-gallon tanks in a recirculating aquarium system with artificial seawater for a one-month acclimation period. Acclimation conditions were based on established husbandry protocols (Rotjan et al. 2019); temperatures were maintained at 21°C, with a 12-hour light/dark cycle. Colonies were fed 3 times per week during acclimation.

### Immune Challenge

Following the acclimation period, colonies of *A. poculata* were exposed to an experimental immune challenge consisting of either heat-inactivated *Vibrio coralliilyticus* (1 × 10^7^ CFU in 100uL of buffer) or a buffer control (filter-sterilized artificial sea water (FSW)) via injection (sample sizes in **Table 1**). Cultures of the known coral pathogen *V. coralliilyticus* (Pollock et al. 2010) were obtained from David Nelson at the University of Rhode Island, *V. coralliilyticus* (strain RE22Sm). Cultures of *V. coralliilyticus* were revived and grown overnight in mYP30 (0.5% yeast extract, 0.1% peptone, 3% marine salts at 27C (Karim et al. 2013). Cultures were then diluted to a final concentration of 1 × 10^8^ CFU, resuspending in FSW, and heat-inactivated by heating to 90°C for 5 minutes (heat inactivation was confirmed via plate streaking)..

**Table 1.** Full sampling scheme used in the final experiment.

|  | *Brown* | *White* |
| --- | --- | --- |
| *Control* | 15 | 4 |
| *Immune Challenge* | 11 | 7 |

Colonies of *A. poculata* were injected with 100uL of either heat-inactivated bacteria or FSW using a PrecisionGlide Needle (BD). The total inoculant (100uL) was divided across 4 injection locations roughly spread across the surface of the coral colonies. Following injection, samples were briefly held in individual containers to allow for inoculate absorption, before being placed back into isolated aquaria for the duration of the experiment. Colonies were randomly divided across 4 aquaria for the incubation period (two per treatment group). Following an incubation period of four hours, samples were removed and subsampled for transcriptomic analyses. Samples preserved for transcriptomic analyses were stored in 1mL of RNAlater (Sigma-Aldrich) for 24 hours at 4°C, then stored in -20°C. The remaining colony was flash frozen in liquid nitrogen for symbiont density analyses.

### RNA Extraction

RNA was extracted from sub-samples preserved in RNA-later using the RNAqueous extraction kit (Ambion by LifeTechnologies) following manufacturer’s protocols, with the addition of a bead beating step prior to lysis to separate coral tissue from the skeleton. The sample was added to a tube with a approximately 0.25mL of 0.5mm glass beads and 500uL lysis buffer and beat on a bead beater (Fisherbrand Bead Mill 24 Homogenizer) for 1 minute (Wuitchik et al. 2021). Following extraction, samples were qualitatively assessed using the Take3 plate on the Cytation 1 cell imaging multi-mode reader with Gen5 software (BioTek). Samples which met established quality guidelines set by the sequencing facility were sent to University of Texas Genomic Sequencing and Analysis Facility for TagSeq library prep and sequencing on NovaSeq with single end, 100 base pair reads.

### Symbiont Density Estimation

Symbiont density was estimated following established protocols (Changsut et al. 2022). Briefly, tissue was removed from a fixed surface area on the flattest portion of the coral and homogenized to generate a tissue slurry. The slurry was then serially washed and centrifuged to isolate photosymbionts, which were stored in 500μL 0.01% SDS in deionized water at 4°C. Symbiodiniaceae density was estimated from this sample following standard protocols (Changsut et al. 2022). Symbiodiniaceae counts were repeated in triplicate and averaged to calculate symbiont density/tissue area.

### Transcriptome Assembly and Annotation

A *de novo* reference transcriptome was assembled from a random subset of 10 colonies. The individuals used in the assembly included 2 brown control, 3 brown infected, 2 white control, and 3 white infected colonies. Novogene conducted mRNA library prep and sequencing on an Illumina NovoSeq 6000 with paired end 150 base pair reads. The resulting sequences were quality assessed using FastQC (Andrews 2010). The sequences were then trimmed to remove adapter sequences and quality filtered to remove sequences with an average Phred score <20 using cutadapt v2.6 (Martin 2011). Since not all sequences contained adapter sequences, the untrimmed output was concatenated to the trimmed output to reduce data loss. Host coral and algal reads were separated using BBsplit v38.84 under default parameters (Bushnell 2014). Reads were mapped against the *B. psygmophilum* reference transcriptome (Parkinson et al. 2016) and those that did not map to the reference were sorted into a new file of coral only reads.

A host-only reference transcriptome was assembled using Trinity v2.8.5 (Grabherr et al. 2011) with the following default parameters. TransDecoder v5.7.1 was used to extract coding sequences from the assembly (Haas and Papanicolaou 2016). These coding sequences were then clustered by similarity using cd-hit (Li and Godzik 2006) to further reduce the size of the assembly. The assembly was then filtered using standard Trinity scripts to retain only the longest isoform for each transcript. Finally, to reduce any missed symbiont contamination, the resulting transcriptome was compared against a custom database of Symbiodiniaceae protein sequences from the following species for which robust protein databases were available: *S. trinacnidorum* (González-Pech et al. 2021), *S. fitti* (Reich et al. 2021), *S. goreaui* (Chen et al. 2022), and *S. kawagutii* (Lin et al. 2015) following established protocols (Stankiewicz et al. 2023). Any sequence with at least 80% identity and 100bp matched were filtered from the transcriptome as symbiont contamination (∼500 sequences removed). Quality assessment of the final assembly and intermediate steps was done using transcript counts, N50s, and benchmarking universal single-copy orthologs (BUSCO) v5.5.0 (Simao et al. 2015) to assess completeness. The resulting assembly was annotated via comparison to the reviewed UniProtKB database (blastx). A maximum of five hits per transcriptomic sequence were returned, with an e-value cut off of 1E-5. Results were then filtered to retain only the hit with the highest e-vlaue for each transcript.

### Read Alignment & Differential Expression Analyses

TagSeq samples were processed and aligned to the generated *de novo* assembly following the established pipelines (Meyer et al. 2011). First, reads were demultiplexed, trimmed, and quality assessed using a combination of custom Perl scripts and cutadapt (Martin 2011). Resulting reads were then mapped to the generated reference transcriptome using Bowtie2 (Langmead and Salzberg 2012) to generate a read count matrix. Differential expression analyses were conducted in R using the package DESeq2 (Love et al. 2014). To increase statistical power and reduce effects of lowly expressed transcripts prior to differential expression analysis, the resultant read count matrix was filtered to retain only those transcripts with an average read count of 3. First, we identified genes which were differentially expressed as a result of immune challenge or symbiont density using the model: gene expression ∼ immune challenge + symbiont density. Then, we tested for genes which responded differentially to immune challenge as a result of variation in symbiont density using the model: gene expression ∼ immune challenge*symbiont density. Differentially expressed genes involved in immunity were identified based on biological process gene ontology terms using a search for the keywords: “immun”, “defense”, “inflamm”, and “phagocy”. Classification as a potential immune gene was then verified by literature review. We assessed enrichment of biological process gene ontology (GO) terms within each of our sets of differentially expressed genes using GOMWU (Wright et al. 2015). To perform GOMWU analyses, we using signed log adjusted p-values, and the annotations generated for the *de novo* transcriptome. Significant genes and GO terms were identified as those with an adjusted p-value < 0.1 (10% FDR).

## RESULTS

### *De novo* Transcriptome Assembly and Read Alignment

A total of 219,350,638 paired reads were assembled into a raw assembly of 973,377 transcripts. Following the last symbiont decontamination step, we obtained a final reference transcriptome with 158,336 transcripts and an N50 of 671. The BUSCO scores for the final assembly were 719 complete (378 single-copy, 341 duplicate-copy, total = 75.3%), 159 fragmented (16.7%), and 76 missing (8.0%). Comparison to the UniProtKB database resulted in a total of 75,592 annotated sequences (47.7%). Individual sample mapping rates to the resultant *de novo* transcriptome ranged from 18-35%. After filtering, a total of 11,487 transcripts passed filtering steps and were analyzed for differential expression, with a final total reads per sample average of 424,527 (range of 213,393-659,619).

### Effects of Immune Challenge

A total of 272 transcripts were differentially expressed between control and immune challenge corals (*padj* < 0.1; 10% FDR; **Supplementary File 1**). Using gene ontology enrichment analysis to examine broad patterns of biological processes, we identified twelve biological process GO terms that were significantly differentially expressed as a result of immune challenge (**Supplementary Figure 1; Supplementary File 2**). The majority of these (8/12) were involved in ciliary, axoneme, or microtubule processes, all of which were negatively enriched in immune-challenged colonies. When considering genes and processes involved in immune response specifically, we identified 12 potential immune transcripts and 2 gene ontology terms. Of the 12 differentially expressed immune transcripts, all but two were expressed higher in corals from the immune challenge group (**Figure 1; Supplementary File 3**). Significantly differentially expressed immune genes included essential coral innate immunity transcription factor, NFKB (Williams et al. 2018; Williams and Gilmore 2020). Both immune biological process GO terms were significantly positively enriched: *negative regulation of immune effector process* and *antimicrobial humoral response* (**Supplementary Figure 1**).

**Figure 1.**
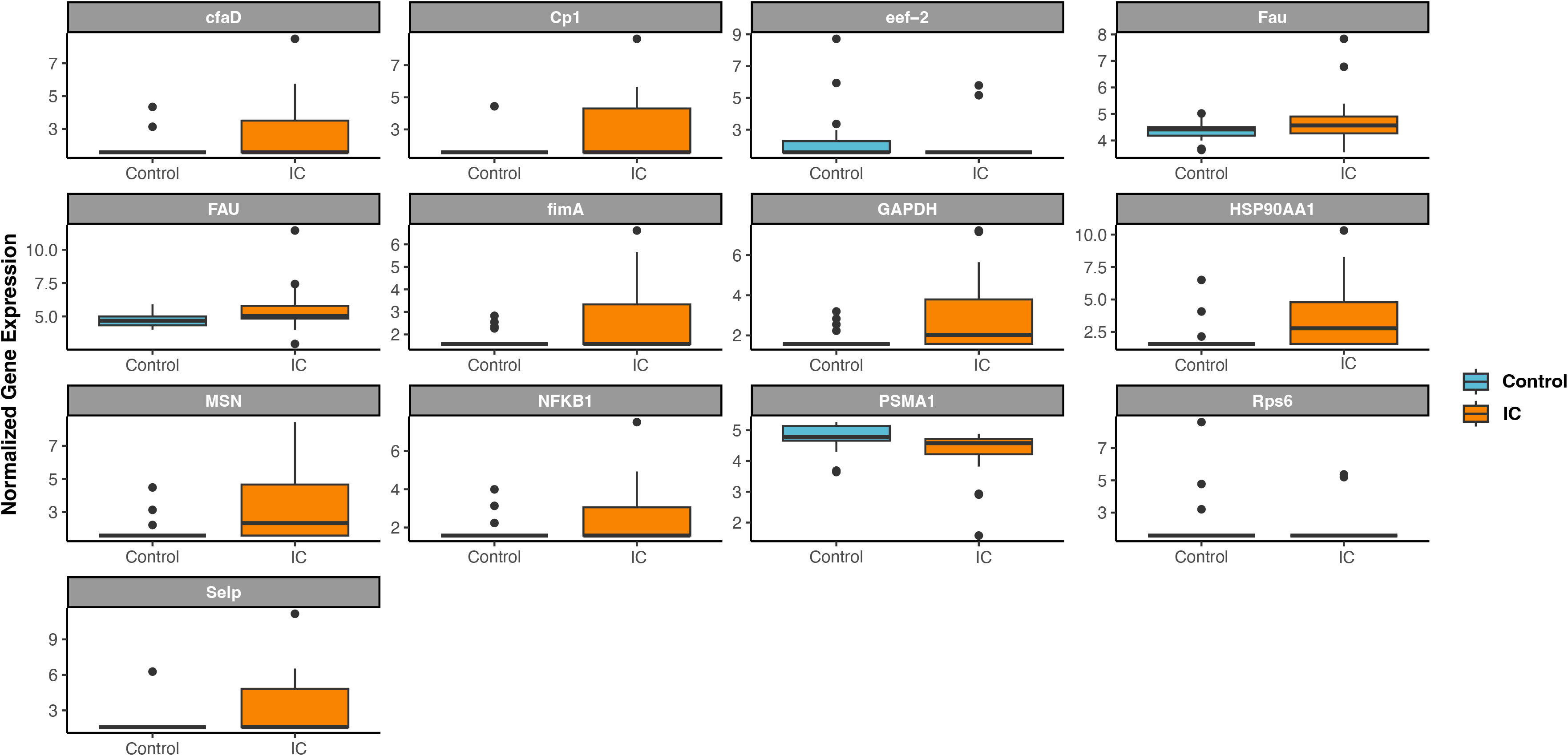
Box plot displaying normalized gene expression of immune genes which were significantly differentially expressed as a result of the main effect of immune challenge (*padj* < 0.1; 10% FDR). Boxes are colored based on treatment; IC = immune challenge. Gene names are displayed for each respective graph.

### Effects of Variation in Symbiont Density

Variation in symbiont density was linked to differential expression of 71 genes, of which 5 were identified as putative immune genes (*padj* < 0.1; 10% FDR). All five identified immune genes were significantly positively correlated with symbiont density (**Figure 2; Supplementary Files 1&3**). Only four biological process GO terms were significantly enriched as a result of symbiont density (**Supplementary Figure 2; Supplementary File 3**). This included *carbohydrate derived catabolic process*, which was negatively enriched (i.e. negatively related to symbiont density). In contrast, *amide metabolic process* was positively enriched (i.e. positively related to symbiont density).

**Figure 2.**
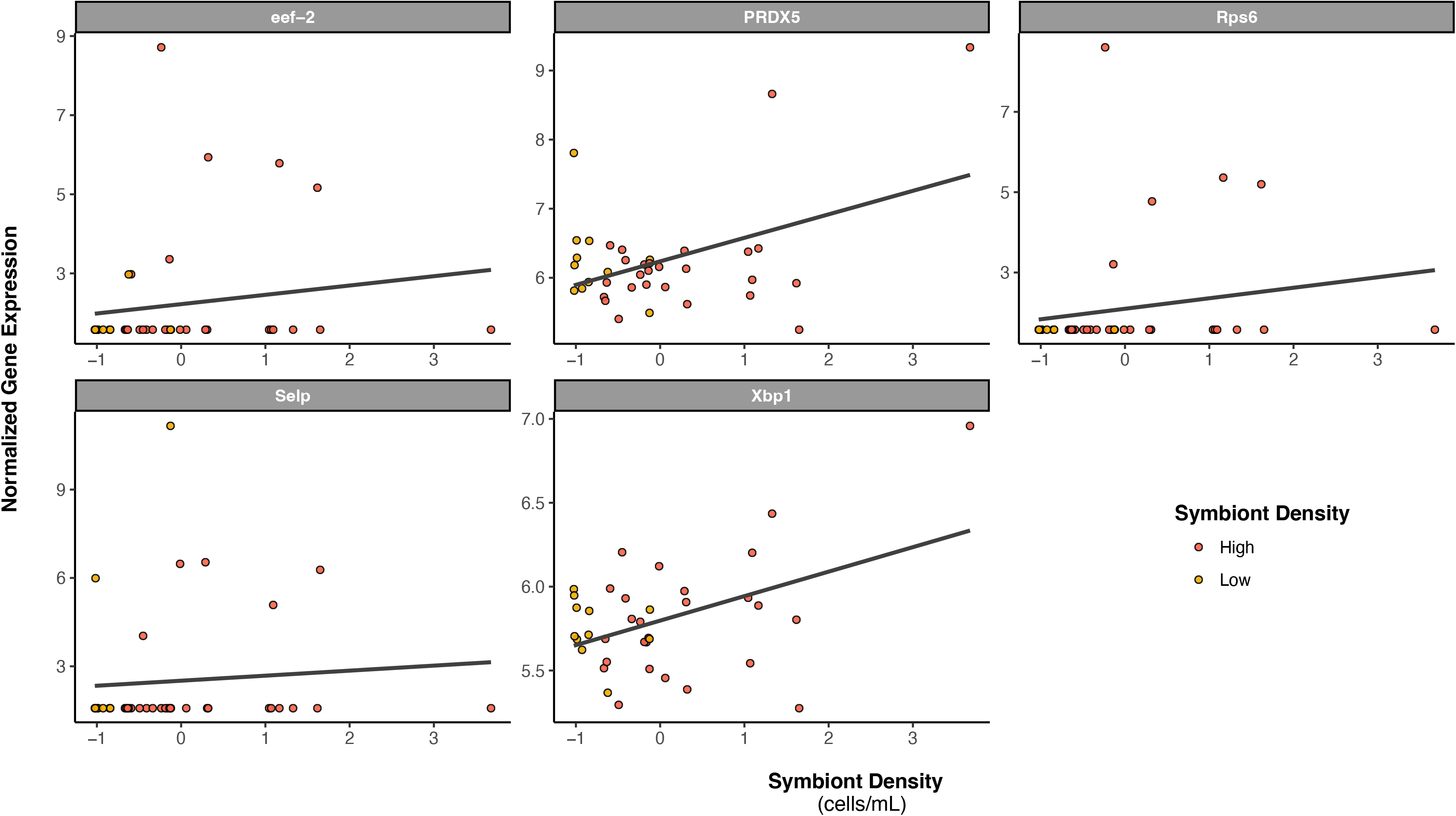
Scatter plot displaying association between normalized gene expression and symbiont density (values scaled and centered) for all significantly differentially expressed immune genes (*padj* < 0.1; 10% FDR). Points are colored based on initial visual colony classification: brown/high symbiont density or white/low symbiont density. Trendlines represent linear model fit of the association between gene expression and symbiont density for all data points. Gene names are displayed in each graph header.

### Variation in Response to Immune Challenge as a Result of Symbiont Density

Finally, 69 genes were identified as differentially expressed by our interaction model (symbiont density*immune challenge effect), meaning that these genes respond differentially to immune challenge as a function of symbiont density. These genes included four putative immune genes, all but one of which (Selp) were expressed higher in response to immune challenge in high symbiont density corals compared to low symbiont density corals (**Fig 3; Supplementary Files 1&3**). Forty-one biological process GO terms were differentially enriched reflective of differences in response to immune challenge due to variation in symbiont density. The majority of these terms (28) were negatively enriched, meaning the processes are more highly induced in response to immune challenge by low symbiont density corals. Of these 28 terms, 13 were involved in ciliary action or related processes (**Fig 4; Supplementary File 2**). In contrast, only 14 terms were positively enriched, of which nine were associated with metabolism/catabolism. A tenth term, *humoral immune response*, was also positively enriched (**Fig 4; Supplementary File 2**).

**Figure 3.**
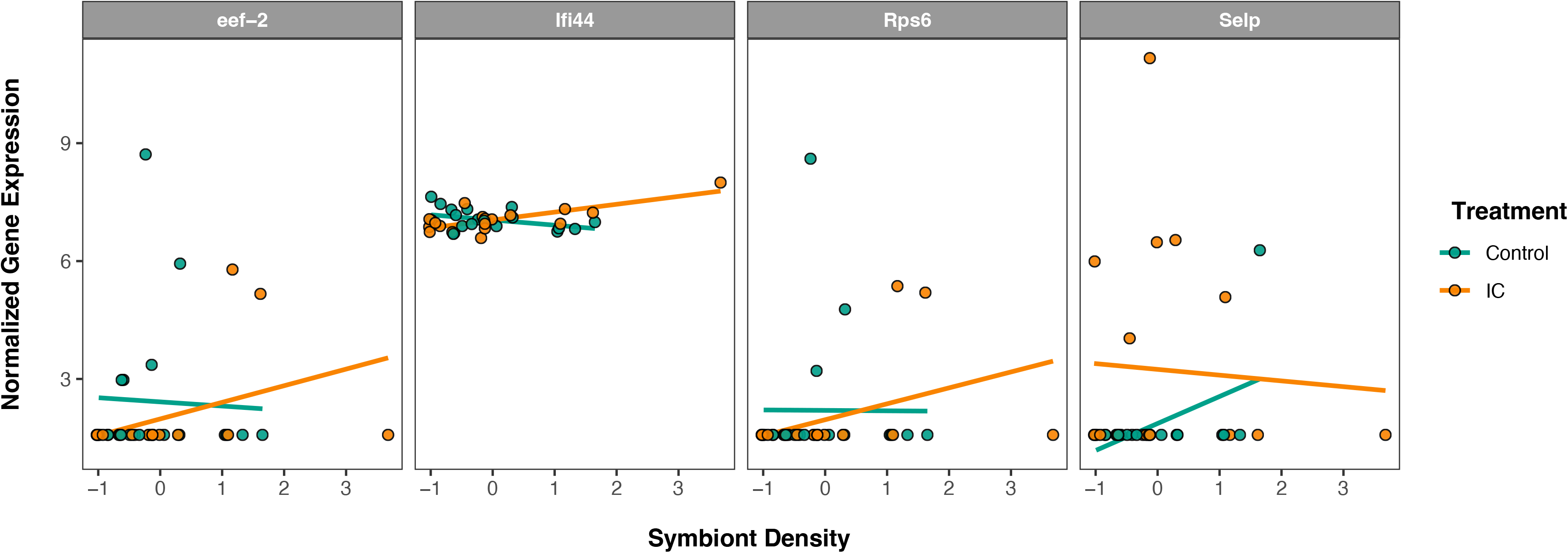
Scatter plot displaying association between normalized gene expression and symbiont density (values scaled and centered) for all immune genes significantly differentially expressed as a result of the interaction between symbiont density and immune challenge (*padj* < 0.1; 10% FDR). Points and trendlines are colored based on treatment group: control or immune challenge (IC). Trendlines represent linear model fit of the association between gene expression and symbiont density within each group (control or IC). Gene names are displayed in each graph header.

**Figure 4.**
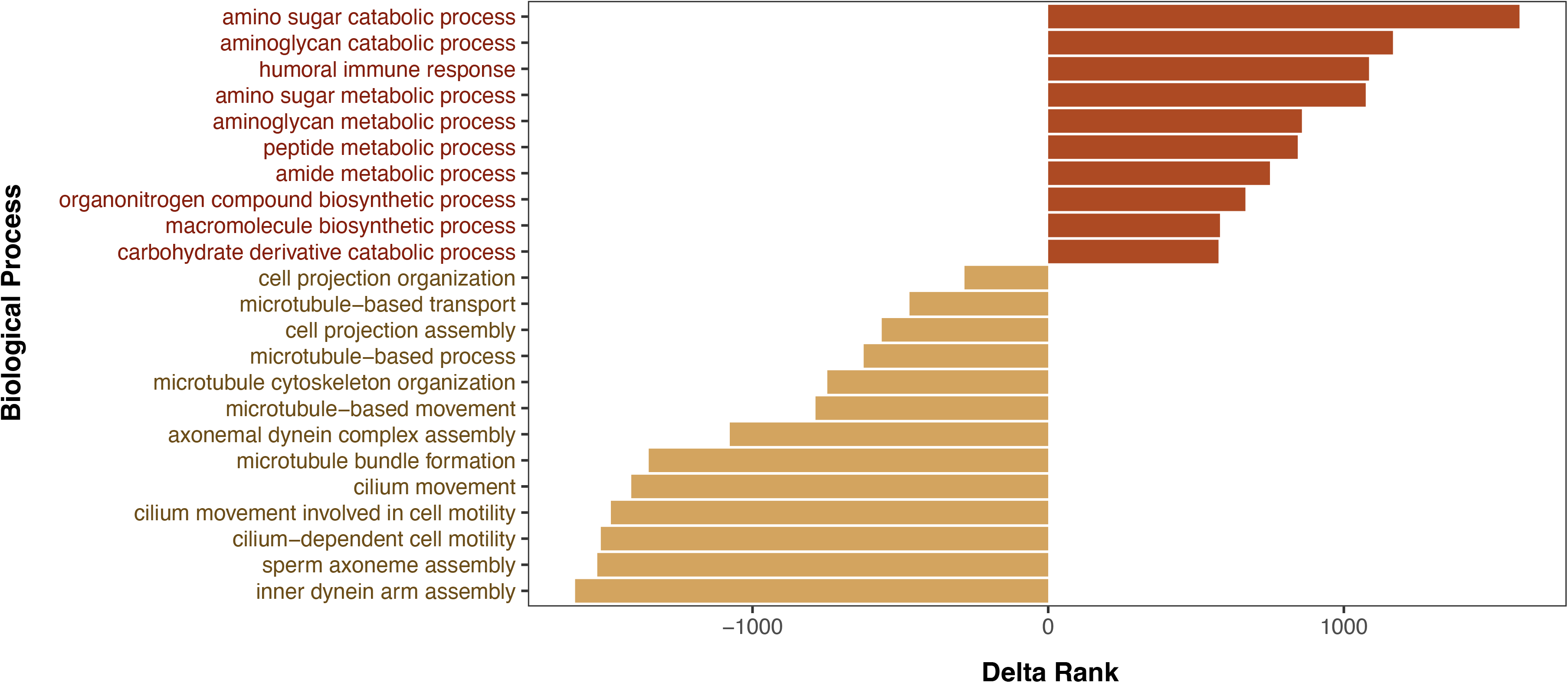
Bar plot representing selected significantly enriched gene ontology terms related to the interaction term symbiont density*immune challenge. Terms displayed are those involved in immune processes, ciliary action (or related processes), and energetic processes. Bar length represents term delta rank, a measure of the magnitude of enrichment. Terms with negative delta ranks characterize the unique response of low symbiont density corals to immune challenge (colored in gold), while terms with positive delta ranks characterize the unique response of high symbiont density corals to immune challenge (colored in red).

## DISCUSSION

Here we leveraged the facultatively symbiotic temperate coral species, *Astrangia poculata*, to investigate impacts of variation in symbiont density on constitutive and induced immune response in cnidarians. Previous studies across a range of scleractinian coral systems have revealed contrasting impacts of symbiont density on host immunity (Changsut et al. 2022; Fuess et al. 2020b; Rivera and Davies 2021; Villafranca et al. 2023). Prior studies of obligately symbiotic corals have indicated negative associations between symbiont density and immunity (Fuess et al. 2020b; Mansfield and Gilmore 2019; Rivera and Davies 2021), while recent work in *A. poculata* have indicated a positive association between symbiont density and a limited suite of constitutive immune metrics (Changsut et al. 2022; Harman et al. 2022; Villafranca et al. 2023). Here we use experimental immune challenges and transcriptomic analyses to comprehensively investigate differences in constitutive immunity between corals of variable symbiont density and to investigate effects of symbiont density variation on induced responses. Our results confirm the findings in previous *Astrangia* studies and highlight important biological processes which may modulate the pathogen response in the facultatively symbiotic coral, *A. poculata*.

### *Astrangia poculata* displays a robust response to immune challenge

Here we present the first transcriptomic data characterizing the response of a facultatively symbiotic coral, *A. poculata* to pathogenic stimuli. Our findings suggest that *A. poculata* mounts an immune response that is similar to those described in other cnidarian systems. Specifically, induction of NFkB is characteristic of immune response across multiple cnidarian species (Fuess et al. 2017; Ocampo and Cadavid Gutierrez 2014; Williams et al. 2018; Williams and Gilmore 2020). Additionally, we see induction of antimicrobial humoral immune response processes, characterized largely by upregulation of the TLR and NFkB related gene, deleted in malignant brain tumor protein 1 (DMBT1; (Bikker et al. 2002; Fuess et al. 2016; Wright et al. 2017). DMBT1 has frequently been described as an essential component of transcriptomic stony coral immune response and has been linked to antimicrobial production in some species (Fuess et al. 2016; Wright et al. 2015). Combined, our differential expression and gene ontology enrichment analyses suggest a strong role of TLR-induced NFkB signaling as part of the initial *A. poculata* immune response, consistent with a large number of studies from tropical stony corals (Connelly et al. 2020; Palmer and Traylor-Knowles 2012; van de Water et al. 2015; Williams et al. 2018; Williams and Gilmore 2020).

We also document downregulation of ciliary action in the initial immune response. This downregulation is in contrast to numerous previous studies which have highlighted the importance of ciliary action of cnidarian defense response against both pathogenic and other disturbances (Fuess et al. 2020a). Ciliary action falls in the category of primary immunity or preventative mechanisms of defense, as its primary role is to prevent bacterial infection altogether (Stannard and O’Callaghan 2006). Several studies have illustrated a link between ciliary movement & biogenesis and stress condition such as sediment stress (Erftemeijer et al. 2012) as well as the coral infection response (Gavish et al. 2021). Indeed, ciliary action is essential for the regulation and maintenance of the coral’s surface mucosal layer (Brown and Bythell 2005), which is often regarded as the coral’s first layer of defense (Bakshani et al. 2018). Mucopolysaccharides associated with the surface mucus layer can trap or repel bacteria (Rublee et al. 1980). Apical cilia provide water flow to sweep mucus and other trapped particles like bacteria off the surface of the coral (Mullen et al. 2004). The lack of ciliary action in *A. poculata*’s initial immune response may be linked to environmental differences; unlike tropicals, *Astrangia* is notable for its unique ability to withstand heavy sedimentation, often associated with decreased ciliary action (Peters and Pilson 1985). Alternatively, this pattern may be linked to the mechanisms of immune stimulation used (i.e., injection). Comparison of the responses documented here to those associated with live pathogenic challenge will be essential to disentangle the ultimate biological mechanisms driving the observed patterns. Regardless, taken with the observed patterns of upregulation in NFkB, it can be assumed that, generally, *A. poculata* responds quickly with cellular, rather than preventative, defense mechanisms in response to pathogenic stimuli.

### Symbiont density is positively associated with immune gene expression

The transcriptomic data provides the first comprehensive description of the response of *A. poculata* to pathogenic challenge. Additionally, the dataset is consistent with recent findings of positive associations between symbiont density and immune gene expression in *Astrangia*. These findings include expression of peroxidoxin (PRDX5), a potent antioxidant gene (Walbrecq et al. 2015), linked to redox homeostasis and stress response (Levy et al. 2016; Polato et al. 2013). Previous work has found PRDX5 to be linked to symbiosis establishment in tropical corals (Mohamed et al. 2020; Mohamed et al. 2016) as well as general stress and bleaching responses (Levy et al. 2016; Polato et al. 2013). Notably, this finding is also in agreement with previous work documenting positive associations between symbiont density and phenotypic antioxidant activity (catalase) in *A. poculata* (Changsut et al. 2022). In addition to PRDX5, we document a positive association between symbiont density and XBP1, a protein involved in the unfolded protein response (UPR) in other invertebrates (Richardson et al. 2010). Under mitochondrial stress, the UPR has been documented to induce innate immune responses (Janssens et al. 2014; Pellegrino et al. 2014). Recent studies have not only documented this pathway in cnidarians, but have linked activity of the pathway to immune response as well as temperature and salinity stress (Dimos et al. 2019). Indeed, the UPR is believed to be an important cellular pathway, which may link regulation of symbiosis to immunity in cnidarians (Aguilar et al. 2019; Dimos et al. 2019; Helgoe et al. 2024; Ruiz-Jones and Palumbi 2017). The documentation of this pathway, and its association with symbiont density, in a facultatively symbiotic coral provides new avenues for investigation of these cellular pathways in cnidarians.

### Variation in symbiont density affects response to pathogens

While we observed limited patterns of differential gene expression characterizing variation in response to immune challenge as a result of symbiont density, gene ontology enrichment patterns were much more striking. Interestingly, in contrast with broad patterns of immune response observed, low symbiont density corals displayed unique signatures of response to pathogens characterized by induction of ciliary action and related processes. The preferential activation of such preventative mechanisms may be linked to reduced constitutive immunity observed in low symbiont density corals (Changsut et al. 2022; Harman et al. 2022). One possible explanation is that induction of ciliary action is less energetically costly than other types of immune response, and therefore may be favored by low symbiont density corals which may have lower energetic reserves, though differences in energetic resources as a result of symbiont density in *A. poculata* remain unclear (Trumbauer et al. 2021; Villafranca et al. 2023).

In contrast to low symbiont density corals, the response of high symbiont density corals was marked by induction of several metabolic processes as well as humoral immune response, suggesting a mobilization of resources to fuel defense responses. Immune responses are exceptionally costly processes, necessitating resource investment either from stored resources, or partitioned away from other processes (Lochmiller and Deerenberg 2003; Pinzon et al. 2014; Rolff and Siva-Jothy 2003). The unique patterns of resource mobilization and activation of humoral immune response following immune challenge observed here in high symbiont density corals suggests a fundamental difference in resource availability or use between corals of high and low symbiont density. One likely hypothesis would be that increased symbiont density results in increased resources which may be allocated to costly responses, like immunity. Although previous research suggests a correlation between symbiont density and ability of the host to withstand and recover from various stressors, including starvation, thermal stress, and wounding (Burmester et al. 2018; Burmester et al. 2017; DeFilippo et al. 2016; DiRoberts et al. 2021), research to date has not demonstrated a clear link between symbiont density and energetic budget in *A. poculata* (Trumbauer et al. 2021; Villafranca et al. 2023). We hypothesize that methods used to date to study resource allocation in *A. poculata* have failed to capture the nuances of resource allocation or mobilization associated with immune responses. Therefore, there is a clear need for the inclusion of immune responses in energetic studies of *A. poculata*, and other facultatively symbiotic coral species. This will help to identify mechanisms underlying the complex interactions between symbiont density and coral immune response and resilience to disease.

### Conclusions

In summary, here we present a powerful transcriptomic dataset that expands the understanding not only of the immune response of a facultatively symbiotic coral, but also on the effects of variability in symbiont density on these responses. Our findings suggest that *A. poculata*’s immune response includes molecular pathways also used by well-studied tropical corals, with a few key differences pertaining to ciliary action. Furthermore, our results confirm preliminary studies suggesting a positive association between symbiont density and constitutive immunity in *Astrangia*. We provide compelling evidence that this positive association extends to induced immune responses and is potentially linked to uncharacterized variation in energetic budgets. Our results provide a compelling narrative highlighting the need for more nuanced study of immuno-energetics in *Astrangia poculata* and other facultatively symbiotic species to improve basic understanding of the associations between symbiosis and immunity across the diversity of cnidarians.

## DATA AVAILABILITY

The data and code underlying this article are available on Github at https://github.com/Symbiommunity/Changsut2024.git.

## ACKNOWLEDGEMENTS

This work was supported by Texas State University (Startup Funds to LEF). *Astrangia poculata* specimen collection was done in accordance with the Rhode Island Department of Environmental Management Scientific Collector’s Permit, issued to Roger Williams University (permit #2021-03G).

