## Supplementary figures and images for "Photosymbiont density is correlated with constitutive and induced immunity in the facultatively symbiotic coral, *Astrangia poculata*"

### Supplementary Figure 1

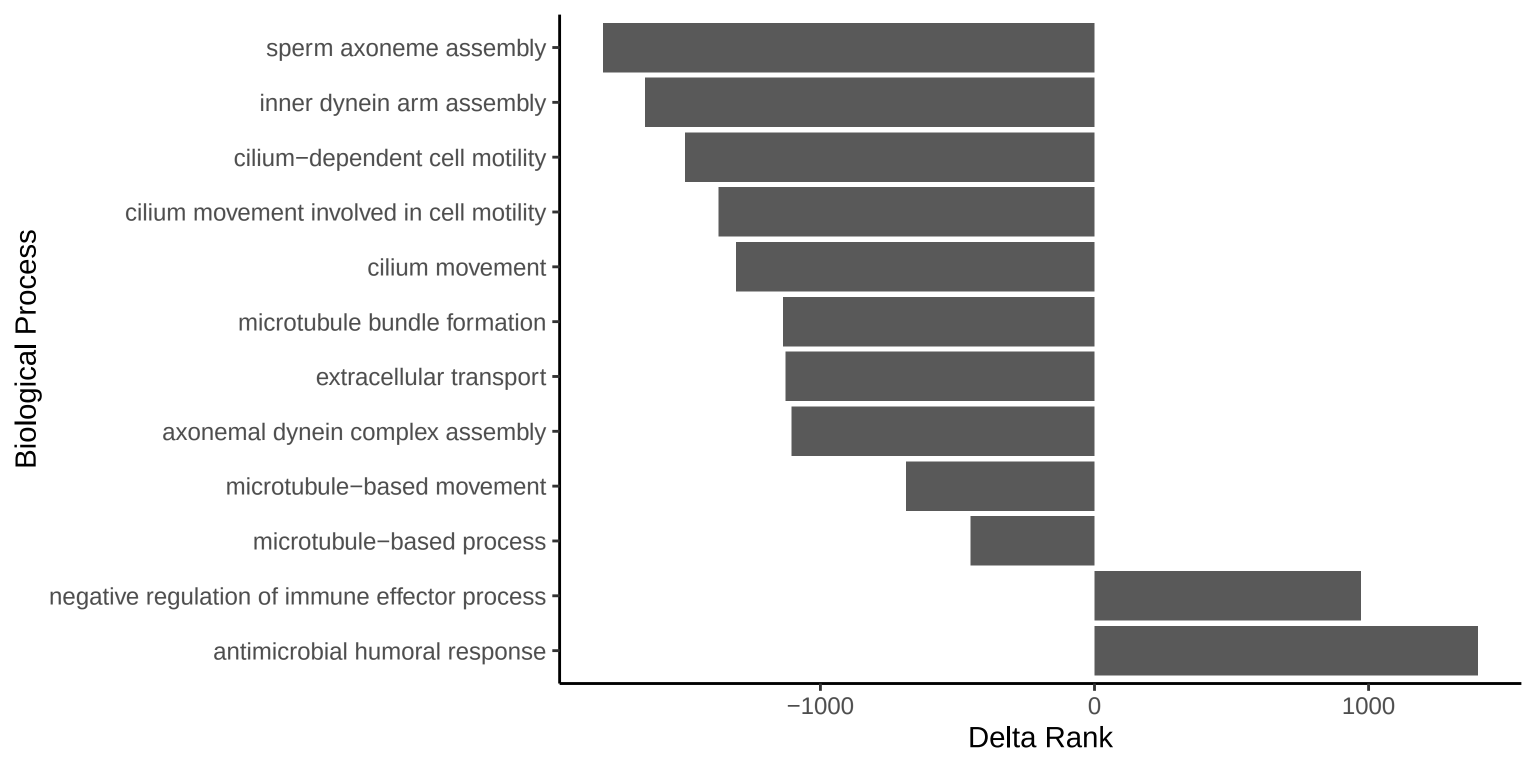

### Supplementary Figure 2

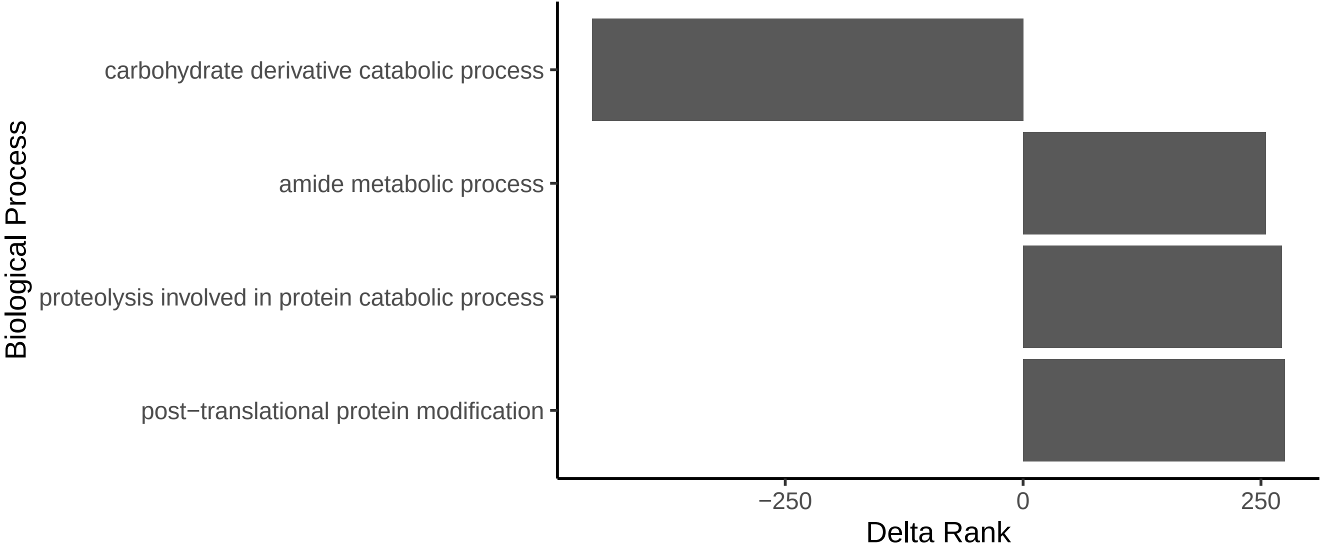
